# L-norepinephrine Induces Community Shift, Oxidative Stress Response, Metabolic Reprogramming, and Virulence Potential in Wastewater Microbiomes

**DOI:** 10.1101/2022.06.28.482069

**Authors:** Amrita Bains, Sanjeev Dahal, Bharat Manna, Mark Lyte, Laurence Yang, Naresh Singhal

## Abstract

Neuroendocrine compounds discharged into wastewater systems represent an emerging challenge at the intersection of human physiology and environmental microbiology. L-norepinephrine (L-NE) L-NE, which has been recognized to potentiate growth of human and animal bacterial pathogens, is discharged in sewage through urine and feces. While extensive pure culture studies have established L-NE’s capacity to modulate bacterial virulence through iron acquisition and quorum sensing pathways, its impact on complex microbial communities, where intricate metabolic networks and interspecies interactions dominate, remains largely unexplored. This knowledge gap is particularly critical as urbanization drives increasing neuroendocrine compound loads in wastewater influents in metropolitan areas. Through parallel treatments of L-NE (1×10□□ to 1×10□□ M), dextrose, and H□O□ in municipal and agricultural wastewater communities, we uncovered sophisticated metabolic and regulatory mechanisms that challenge the conventional understanding of microbial substrate utilization. Despite containing 10-fold less carbon, L-NE treatments achieved superior growth (10□ CFU mL□¹) while maintaining *Pseudomonadaceae*-dominated communities. Targeted metaproteomics revealed coordinated upregulation of oxidative stress genes (*oxyR*, *soxRS*) and antioxidant enzymes, while proteome-constrained metabolic modeling demonstrated distinct pathway modulation in central carbon and nitrogen metabolism. Notably, when compared to dextrose-supplemented controls representing typical carbon substrate utilization, L-NE treatments showed similar taxonomic profiles without preferential enrichment of known pathogenic families. However, L-NE significantly enhanced autoinducer gene (*luxS*, *qseC*) expression, suggesting increased virulence potential through community-level metabolic reprogramming. These findings reveal L-NE as a potent modulator of microbial community dynamics in engineered ecosystems, with important implications for treatment process stability and downstream environmental impacts.

**TOC Figure.**
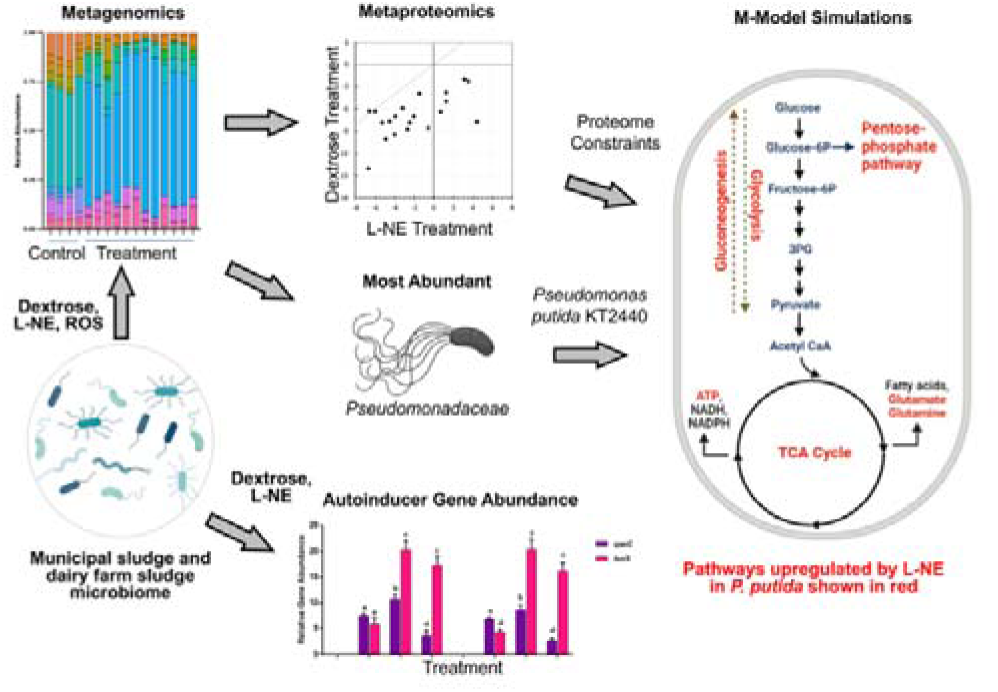

## 1. Introduction

L-norepinephrine (L-NE) and other catecholamines are bioactive compounds that enter wastewater through multiple pathways, including human urine, feces, and plant matter, and have been shown to significantly influence microbial behavior in environmental systems (Freestone et al., 1999; Kinney et al., 2000; Kulma and Szopa, 2007; Lyte, 2016). These neuroendocrine molecules regulate critical bacterial phenotypes, including motility, growth, adhesion, biofilm formation, and virulence through complex cellular mechanisms (Bansal et al., 2007; Chen et al., 2003; Cogan et al., 2007; Gao et al., 2019; Lyte, 2016; Nakano et al., 2007; Sandrini et al., 2014; Sarkodie et al., 2019; Truccollo et al., 2020). At the molecular level, L-NE modulates bacterial gene expression, particularly affecting metabolic processes, DNA repair mechanisms, and ribosomal protein biosynthesis (Gao et al., 2019; Xu et al., 2015). A key aspect of L-NE metabolism is the generation of hydrogen peroxide (H_2_O_2_) and other reactive oxygen species (ROS) (Maggiorani et al., 2017; Neri et al., 2007; Saller et al., 2012), which exhibit dual roles in bacterial physiology: at low concentrations, ROS function as essential signaling molecules promoting growth-related pathways (Bonavita and Laukkanen, 2021; Sies, 2017; Thannickal and Fanburg, 2000), while elevated levels induce oxidative stress, leading to biomolecular damage and cellular death (Imlay, 2013; Sies and Jones, 2020; Yang et al., 2019). In nutrient-limited aerobic environments, catecholamines enhance bacterial growth up to 10-fold across diverse Gram-positive and Gram-negative species (Freestone et al., 1999; Gao et al., 2019; Kinney et al., 2000), primarily through the production of auxiliary siderophores (Boyanova, 2017; Kinney et al., 2000) and autoinducers (Freestone et al., 1999; Lyte et al., 1996; Voigt et al., 2006). These molecular mechanisms have significant implications for bacterial pathogenicity - siderophores enable pathogenic bacteria to compete for limited iron resources, while the autoinducer genes *luxS* and *qseC* regulate virulence expression (Palaniyandi et al., 2013; Rasko et al., 2008). The combined effects of ROS stress (Fones and Preston, 2012; Wan et al., 2017) and enhanced siderophore and autoinducer production create conditions that may promote pathogenesis and increase virulence potential within microbial populations.

L-NE acts as both a signaling molecule and can enhance bacterial iron acquisition by host proteins like transferrin and lactoferrin (Freestone et al., 2000), while simultaneously triggering sophisticated molecular responses across diverse bacterial taxa. In pure culture studies, L-NE has been shown to enhance pathogen survival through iron acquisition (Beasley et al., 2011; Williams et al., 2006) and promote proliferation *via* autoinducer production (Lyte et al., 1996; Voigt et al., 2006). In pathogenic species like *Pseudomonas aeruginosa* and *Vibrio cholerae*, L-NE induces quorum sensing pathways (*las*, *rhl*, *luxS*) and upregulates virulence genes (Halang et al., 2015; Hegde et al., 2009; Xu et al., 2015), while in organisms like *Campylobacter jejuni* and *Salmonella typhimurium*, it enhances motility, adhesion, and biofilm formation through interkingdom signaling mechanisms (Cambronel et al., 2020, 2019). However, despite this extensive characterization in pure cultures, L-NE’s effects on complex microbial communities remain unexplored. This knowledge gap is particularly significant given the potential dual mechanisms of L-NE’s influence: first, as a selective agent through ROS generation, where L-NE metabolism produces hydrogen peroxide and other reactive oxygen species that may provide advantages to organisms with enhanced stress tolerance (Ma et al., 2020; Scardaci et al., 2021); and second, as a signaling molecule that can reprogram microbial metabolism in both pathogenic and non-pathogenic strains through quorum sensing-mediated pathways (Hegde et al., 2009; Xu et al., 2015). The complexity of these interactions in mixed microbial consortia is further amplified by L-NE’s demonstrated ability to generate strain-specific autoinducers (Freestone et al., 1999; Lyte et al., 1996) and influence interspecies competition (Mart’yanov et al., 2021), suggesting potential community-wide effects that cannot be predicted from pure culture studies alone.

Understanding L-NE’s effects on complex microbial communities has become increasingly critical due to its persistent detection in wastewater systems, evidenced by the consistent presence of catecholamine metabolites - vanillylmandelic acid (VMA) and homovanillic acid (HVA) - across 38 Australian wastewater catchments (Pandopulos et al., 2021). HVA represents the primary metabolite derived from dopamine, while VMA forms as one of the main metabolites from epinephrine and norepinephrine (Beals et al., 2022). Metabolite monitoring reveals significant per-capita mass loads of 1.70 ± 52 mg.day^-1^person^-1^ for VMA and 1.94 ± 54 mg.day^-1^person^-1^ for HVA in municipal wastewater, with metropolitan areas showing higher concentrations (1.97 and 2.26 mg.day^-1^person^-1^ respectively) compared to regional sites (1.56 and 1.77 mg.day^-1^person^-1^) (Pandopulos et al., 2021). These metabolite levels indicate substantial neurotransmitter inputs from multiple anthropogenic sources: direct human excretion, pharmaceutical residues, and healthcare facilities. The stability analysis of these metabolites demonstrates their persistence under various wastewater conditions, with both compounds remaining stable for up to 14 days (Pandopulos et al., 2021, 2020). This stability raises particular concerns as conventional biological wastewater treatment processes, while effective for nutrient removal, are not specifically designed to eliminate neuroendocrine compounds or their metabolites. Recent analytical developments using LC-MS/MS have improved detection capabilities for these compounds, demonstrating a strong correlation between metabolite mass loads and catchment population (Pandopulos et al., 2021). Given L-NE’s demonstrated effects on bacterial physiology and the strong linear correlation between its derived metabolites and population size, understanding its influence on microbial community dynamics becomes crucial for wastewater treatment efficacy. This knowledge gap becomes increasingly significant as urbanization intensifies, with metropolitan catchment data showing consistently higher metabolite loads that suggest elevated L-NE presence in more densely populated areas.

In this study, we clarify the mechanistic basis of L-NE’s effects on wastewater microbial communities, their implications for treatment process stability, and downstream environmental impacts. Specifically, we address three fundamental questions regarding L- NE’s influence on wastewater microbial communities: (1) How does L-NE function as both a metabolic substrate and signaling molecule in complex microbial consortia? (2) What regulatory mechanisms govern oxidative stress responses in L-NE-exposed communities? (3) Does L-NE exposure alter community-level virulence traits independent of taxonomic shifts? To investigate these questions, we study microbial communities in wastewater and dairy farm environments using various treatments, including L-NE, to examine the mechanistic impacts on microbial composition and function using advanced molecular techniques like 16S rRNA profiling, metaproteomics, and metabolic modeling.

## 2. Material and methods

### 2.1 Batch experiments

Sludge samples were collected from the settling pond at a dairy farm (FS) and activated sludge from a municipal wastewater treatment plant (MS) in Auckland, New Zealand. The microbial cultures were acclimated by adding 10 mL of sludge to 20 mL of synthetic wastewater containing 400 mg/L-COD of acetate, 60 mg-N/L of ammonium supplemented with 25 mM of HEPES buffer (DI water) and incubated overnight at 37°C on a rotating shaker. Acclimated cultures were centrifuged (4,000 rpm, 10 min) and washed twice with phosphate-buffered saline (PBS). We used a standard curve of A_595_ versus colony-forming units (CFU) to seed 1L bioreactors to 100 CFU mL^-1^ with the two microbial cultures. The final concentrations of bacteria inoculated into bioreactors were determined on tryptic soy agar (TSA) plates using the standard pour-plate technique. Batch experiments were conducted using 1×10^-5^ M, 5×10^-5^ M, and 1×10^-4^ M L-NE in aerated bioreactors to simulate physiologically relevant L-NE levels observed *in vivo* (Lyte et al., 1996). Notably, L-NE bitartrate was used based on a previous report (Lyte and Ernst, 1992). Bioreactors with identical dextrose levels served as controls for easily degradable carbon, noting that dextrose controls had 10-fold higher carbon as compared to L-NE reactors. Two bioreactors seeded with 5, 10, and 50 M H_2_O_2_ added to municipal sludge cultures served as ROS controls. Bioreactors were continuously mixed (100 rpm) and aerated at 22-25°C for 24 h (see Table S1 for conditions). Microbial growth in bioreactors was measured using the standard pour plate technique on tryptic soy agar (TSA) plates at 0, 8, and 24 h. The 24 h samples were analyzed in duplicate for bacterial growth, total abundance of ROS genes *oxyR* and *SoxRS*, autoinducer genes *luxS* and *qseC*, proteins and microbial composition (Table S2). Additional methodological details are provided in Supplementary Information S1.1.

### 2.2 Microbial DNA isolation and taxonomic profiling

Microbial community composition was characterized by amplifying and sequencing bacterial 16S rRNA) gene fragments with the universal 16S PCR forward primer (5’- **TCGTCGGCAGCGTCAGATGTGTATAAGAGACAG**CCTACGGGNGGCWGCAG-3’) and the 16S PCR reverse primer (5’-**GTCTCGTGGGCTCGGAGATGTGTATAAGAGACAG**GACTACHVGGGTATCTAA TCC-3’) following standard protocols (Quast et al., 2013). Nucleotide bases in bold are Illumina overhang adapter sequences for high-throughput sequencing. Amplified PCR products were purified using an AMPure XP beads kit (Beckman Coulter Inc.). DNA concentrations were recorded using a Qubit^®^ dsDNA HS Assay Kit (Life Technologies) and then sequenced using Illumina MiSeq (New Zealand Genomics Ltd., Auckland, New Zealand) using 2-by-300 bp chemistry. Before DNA sequencing, the sequencing provider attached a unique combination of Nextera XT dual indices (Illumina Inc., USA) to the DNA of each sample to allow multiplex sequencing. The resulting paired-end read DNA sequence data were merged and quality-filtered using the USEARCH sequence analysis tool (Edgar, 2013). Data was de-replicated to retain a single copy of each sequence. Sequence data were then checked for chimeric sequences and clustered into groups of operational taxonomic units (OTUs) based on a sequence identity threshold equal to or greater than 97% (hereafter referred to as 97% OTUs) using the clustering pipeline UPARSE (Edgar, 2013) in QIIME v. 1.6.0 (Caporaso et al., 2010). Prokaryote phylotypes were classified according to their corresponding taxonomy by implementing the RDP classifier routine (Wang et al., 2007) in QIIME v. 1.6.0 (Caporaso et al., 2010) to interrogate the Greengenes v13.8 database (McDonald et al., 2012). All chloroplast and mitochondrial DNA sequences were removed. Finally, DNA sequence data were rarefied to a depth of 5,600 randomly selected reads per sample, and two samples per treatment to achieve a standard number of sequencing reads across all samples. All bioinformatics analyses were performed in R version 4.2.1.

### 2.3 Protein extraction and identification

Enzymes for oxidative stress and central carbon metabolism (TCA and nitrogen cycle) were extracted by lysing cells in the dairy farm and municipal sludge cultures. One ml of the culture was pelleted (16,000g, 5 min) at 4°C and washed twice with 50 mM ammonium bicarbonate. The washed pellets were resuspended in 150 µl of 7M urea-thiourea buffer and sonicated for 15 s in three rounds on ice. The extracted proteins were separated from residual cellular material by centrifuging at 16,000 xg for 5 min. The proteins in the supernatant were quantified using the EZQ fluorescence protein assay (EZQ protein quantification kit) as per the manufacturer’s protocol. Briefly, Trypsinization was performed on samples, and the proteins were recovered using HLB solid phase extraction cartridges. Finally, 30 µL of the generated tryptic peptides of the microbial proteins were injected into a SCIEX 6600 triple TOF mass spectrometer (AB Sciex, Australia). The proteins were identified by comparing the peptide sequences against UniProt database (Uniprot.org). The complete methodology is presented under Supplementary Information S1.3.

### 2.4 Targeted metaproteomics data analysis

Two separate pipelines for matching peptides were pursued to identify peptide-associated functions. Briefly, in the first pipeline, the Peptide Search tool (https://www.uniprot.org/peptidesearch/) was used for gene ontology (GO) assignment. In the second pipeline, the Metaproteomics Analysis tool (https://unipept.ugent.be/datasets) in Unipept (Gurdeep Singh et al., 2019) was used to assign GO terms. Then, results for these two pipelines were merged (Supplementary Data SD1), and molecular functions (GO terms) of the microbial communities were analyzed across treatments. First, the peptide intensities for the mapped GO terms were summed and the average of the logs of the total intensities for biological replicates was calculated following data normalization using intensities computed for HDO condition. A two-sided t-test was performed to identify the functions that changed significantly. For the t-test, the null hypothesis assumed that the averages of two independent samples were identical. Scatterplots were utilized to visualize the results of the analyses. The complete methodology is presented under Supplementary Information S1.4.

### 2.5 M-model simulations of archetypes

Archetypal analysis (AA) finds “archetypes”, representing points within the multivariate data whose convex combination can well represent the data set (Cutler and Breiman, 1994). We performed archetypal analyses as described in (Yang et al., 2019). The archetypal analysis was performed using the *py_pcha* tool, which is a Python implementation of principal convex hull analysis (https://github.com/ulfaslak/py_pcha), to analyze the matrix to determine the pathway differences between dextrose and L-NE growth conditions. Because *Pseudomonadaceae* was the most abundant family of the microbiome, a modified model of *Pseudomonas putida* (*i*JN1463) (Supplementary Information S1.5) was simulated for a range of growth rates to create a flux distribution matrix (Nogales et al., 2020). For the simulation, most fluxes were chosen based on the previous studies on M-models and their concentration in the medium (Bartell et al., 2017). The oxygen uptake rate constraint was approximated using another study (Gomez et al., 2006). The fluxes for acetate, ammonium, and methanol were arbitrarily chosen based on their concentration in the experimental medium. The uptake fluxes for L-NE and dextrose for their respective medium were constrained at 6 mmol gDW^-1^ hr^-1^ which was the default uptake rate for dextrose set in the model. The complete methodology is presented under Supplementary Information S1.5.

## 3. Results and Discussion

### 3.1 L-NE Selects for *Pseudomonadaceae*-dominated communities

To investigate L-NE’s influence on microbial community dynamics, we exposed microbial consortia from dairy farm sludge (FS) and municipal sludge (MS) to physiologically relevant L-NE concentrations (1×10-5, 5×10-5, and 1×10-4 M), previously established to affect bacterial behavior *in vivo* (Lyte et al., 1996). Parallel cultures with equivalent molarities of dextrose served as carbon source controls, while H_2_O_2_ treatments (5 μM, 10 μM, and 50 μM) provided oxidative stress controls. L-NE significantly enhanced microbial growth in both sludge communities, with the most pronounced effect observed at 5×10^-5^ M L-NE, reaching 10^8^ CFU mL^-1^. Lower (1×10^-5^ M) and higher (1×10^-4^ M) L-NE concentrations yielded moderate growth increases to 10^6^ and 10^7^ CFU mL^-1^, respectively (Figure S1). Notably, despite containing approximately 10-fold higher carbon content, dextrose treatments showed markedly lower growth stimulation, reaching only 10^5^ CFU mL^-1^ across all tested concentrations (Table S3). This enhanced growth response to L-NE persisted throughout the experimental period, with sustained higher cell densities at both 8 and 24 hours compared to dextrose controls.

Correlation analysis of bacterial family abundances revealed distinct community responses between treatments (Figure 1). Using high dissolved oxygen (HDO, 8 mg/L) conditions as a normalization baseline, we observed strong linear correlations between dextrose (5×10^-5^ M) and L-NE treatments in municipal sludge communities (R^2^ = 0.92, 0.90, and 0.89 for 1×10^-5^, 5×10^-5^, and 1×10^-4^ M L-NE, respectively). Notably, the 5×10^-5^ M dextrose concentration was selected as the primary comparison point based on its optimal growth stimulation among dextrose treatments (Table S3). The dairy farm sludge exhibited similar but progressively decreasing correlations (R^2^ = 0.84, 0.67, and 0.48) with increasing L-NE concentrations, suggesting source-dependent community responses. Importantly, comparisons between dextrose and H_2_O_2_ treatments showed minimal correlation (R^2^ = 0, 0.01, and 0.14 for 5, 10, and 50 μM H_2_O_2_, respectively), indicating distinct community responses to oxidative stress versus carbon source availability.

**Figure 1:**
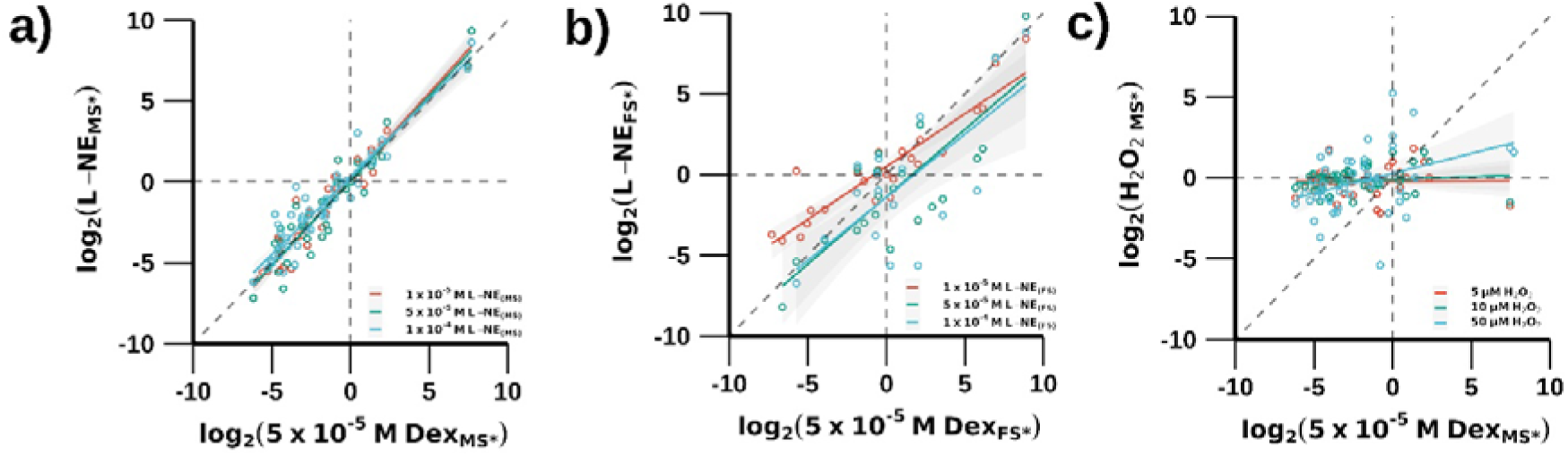
Correlation graphs comparing the effect of L-NE and ROS with dextrose on common microbial family’s abundances in the municipal (MS) and dairy farm (FS) sludge cultures. Effects of L-NE and dextrose on microbial abundances are shown in (a) MS and (b) dairy FS. The effects of H_2_O_2_, vs dextrose is presented in (c). The standard error values of the fitted line are provided in Table S4. (Note: *represents data scaled to observed abundances for the treatment with the corresponding values for HDO_(MS)_).

The family-level abundance profiles in these oxidative stress conditions aligned more closely with ROS-treated samples than with dextrose treatments. Alpha-diversity analysis revealed higher microbial community diversity in both HDO control and H_2_O_2_-treated systems compared to L-NE treatments (Figure S2), with L-NE exposure triggering dramatic shifts in community composition relative to both HDO and H_2_O_2_ conditions. Taxonomic profiling revealed distinct community structures across treatments (Figure S3). HDO control and H_2_O_2_-treated systems shared similar taxa profiles, dominated by *Chitinophagaceae*, *Saprospiraceae*, *Cytophagaceae*, *Flavobacteriaceae*, *Xanthomonadaceae*, and *Rhodocyclaceae*. In contrast, L-NE and dextrose treatments induced a shift toward *Pseudomonadaceae* dominance, accompanied by other prominent families, including *Moraxellaceae*, *Comamonadaceae*, *Weeksellaceae*, *Sphingobacteriaceae*, and *Methylophilaceae*.

Microbial community dynamics in wastewater treatment systems are fundamentally shaped by environmental and operational parameters. Our findings demonstrate distinct patterns of community restructuring in response to L-NE exposure as a known biomass growth stimulant *in vivo* (Freestone et al., 1999). While both L-NE and dextrose enhanced bacterial growth in MS and FS bioreactors, the enrichment patterns revealed source-dependent variations in community response. Notably, *Pseudomonadaceae* emerged as the dominant family across both MS and FS cultures under L-NE and dextrose treatments. The stronger correlation of common family abundances in MS compared to FS treatments underscores the influence of source environment on community restructuring potential. Although L-NE and dextrose treatments yielded similar community compositions, they induced distinct variations in relative abundances. Importantly, potential pathogenic families, e.g., *Enterobacteriaceae*, *Moraxellaceae*, and *Neisseriaceae,* remained at low abundance across all experimental conditions, suggesting that L-NE exposure did not preferentially enrich pathogenic taxa in these wastewater communities.

### 3.2 L-NE Triggers Distinct Oxidative Stress Responses

Given the similar community compositions but different growth patterns between L-NE and dextrose treatments and their distinct profiles from H_2_O_2_ treatments, we investigated underlying oxidative stress response mechanisms. Using the 16S housekeeping gene as a reference, we quantified the abundance of two key oxidative stress response genes, *soxRS* and *oxyR* (Figure S4). Under HDO and dextrose conditions, both genes maintained low abundance. However, *oxyR* abundance increased significantly in response to 50 µM H_2_O_2_ exposure, providing a reference point for oxidative stress response. All L-NE concentrations (1×10^-5^ M, 5×10^-5^ M, and 1×10^-4^ M) induced elevated *oxyR* abundance, with 5×10^-5^ M eliciting the strongest response. The comparable *oxyR* levels between 5×10^-5^ M L-NE and 50 µM H_2_O_2_ treatments suggested equivalent oxidative stress intensity, though notably, *soxRS* expression remained consistently lower than *oxyR* across all L-NE treatments.

While L-NE and dextrose treatments resulted in similar community compositions dominated by *Pseudomonadaceae*, they exhibited different oxidative stress responses. HDO and H_2_O_2_ conditions promoted higher microbial diversity (Shannon index) compared to L-NE/dextrose treatments while showing reduced correlation with dextrose-associated family abundances. The decreased microbial family abundance under HDO and H_2_O_2_ treatments reflected their distinct stress conditions compared to L-NE/dextrose-supplemented systems. Interestingly, while dextrose treatments maintained similar microbial abundances to L-NE, they showed minimal *oxyR* induction, suggesting fundamentally different community response mechanisms. The molecular basis for these observations lies in L-NE’s complex interactions with cellular metabolism. In L-NE-exposed cultures, intracellular ROS generation occurs through dual mechanisms: constant aeration and oxidative deamination of L-NE by oxidase in the presence of oxygen (Buhlman, 2016; Lyte and Freestone, 2010; Neri et al., 2007). These ROS-generating pathways create selective pressure through both direct toxic effects and secondary signaling functions (Dickinson and Chang, 2011). Bacterial adaptation to these conditions involves sophisticated regulatory networks centered on *soxRS* and *oxyR* activation, controlling the expression of key ROS-scavenging enzymes - superoxide dismutase, catalase, and thioredoxin reductase (Imlay, 2013). The enhanced potential of these enzymes in L-NE cultures compared to dextrose treatments demonstrates active engagement of oxidative stress response pathways, correlating with elevated O_2_^-.^ and H_2_O_2_ production. Remarkably, despite generating H_2_O_2_ levels comparable to exogenous 50 µM H_2_O_2_ treatment, L-NE-exposed communities exhibited distinct abundance patterns, suggesting successful adaptation to ROS stress. This adaptation indicates that L-NE’s influence extends beyond simple oxidative stress, potentially involving additional signaling mechanisms that enable community resilience while maintaining selective pressure for specific microbial families.

### 3.3 L-NE Reprograms Protein Expression and Stress Tolerance

Having established distinct oxidative stress responses despite similar community compositions between L-NE and dextrose treatments, we performed targeted metaproteomics analysis to investigate underlying molecular mechanisms. Gene ontology (GO) categories were estimated using peptides from the 50 µM H_2_O_2_ from municipal sludge cultures and 5×10^-5^ M dextrose and 5×10^-5^ M L-NE from dairy farm sludge treatments (Figure 2, Supplementary Data 1). Interestingly, the L-NE treatment generated responses that were quite distinct from those of dextrose and H_2_O_2_ treatments. Compared to dextrose, L-NE upregulated most oxidative stress-associated functions except for monooxygenase activity (GO:0004497, six peptides) and ammonia monooxygenase activity (GO:0018597, two peptides) (Figure 2a). Proteins with ammonia monooxygenase activity catalyze the conversion of ammonia to hydroxylamine. Additionally, L-NE upregulated glutamate-ammonia ligase activity (GO:0004356, three peptides), indicating a shift in nitrogen flux towards glutamine production. In terms of nitrogen metabolism, L-NE enhanced functions related to anaerobic respiration, including nitrous oxide reductase (GO:0050304, one peptide) and nitrate reductase (GO:0008940, eight peptides). L-NE also induced ROS detoxification through increased thioredoxin-disulfide reductase, thioredoxin peroxidase, catalase, peroxidase, and oxidoreductase activities. Comparison with H_2_O_2_ treatment (Figure 2b) revealed L-NE-specific upregulation of ATP binding, oxidoreductase, peroxidase, lyase, isomerase, glutamate-ammonia ligase, and N_2_O reductase activities. Notably, thioredoxin-disulfide reductase (GO:0004791, one peptide) and thioredoxin peroxidase (GO:0008379, one peptide) activities, involved in redox balancing and H_2_O_2_ detoxification, showed significant (*p* < 0.05) upregulation. L-NE treatment downregulated several functions, including copper ion, nickel cation and iron ion binding, electron transfer, 4Fe-4S cluster and FAD/FADH_2_ binding, and oxidoreductase, ammonia monooxygenase, and heme binding activities. Both L-NE and H_2_O_2_ treatments showed similar levels of superoxide dismutase and catalase expressions, though overall ROS mitigation appeared enhanced under L-NE conditions.

**Figure 2.**
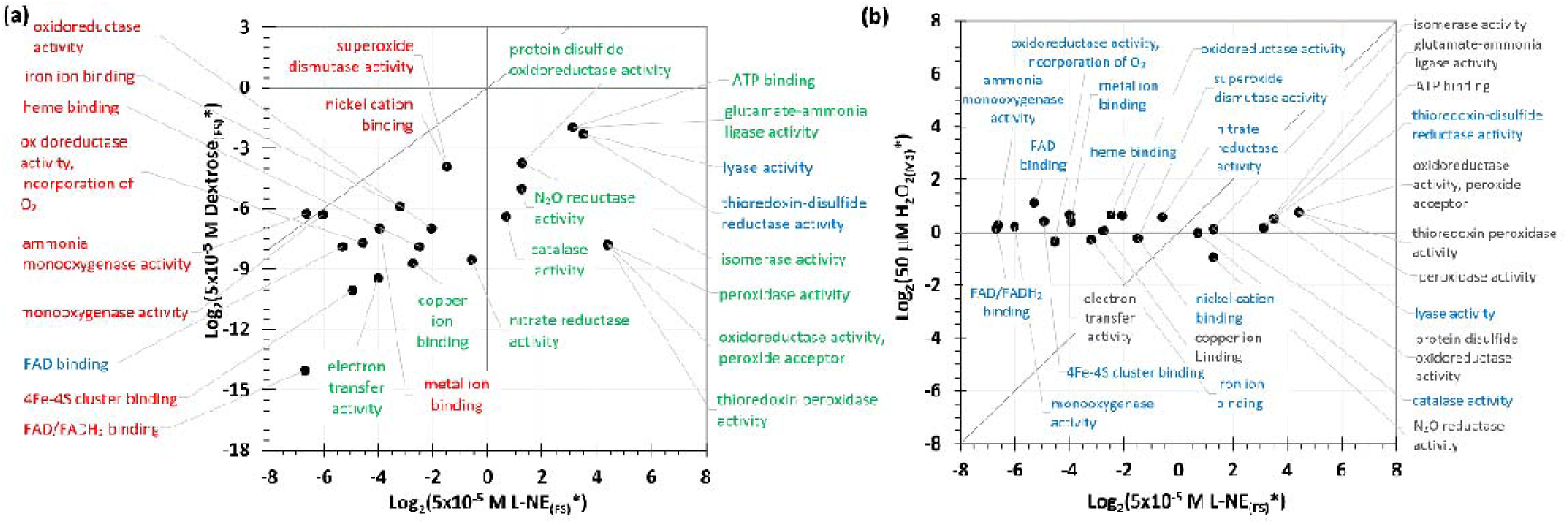
Different organic carbon substrates and oxidative stress conditions cause alterations to protein functions. Plots show: (a) 5×10^-5^ M L-NE vs. 5×10^-5^ M dextrose and (b) 5×10^-5^ M L-NE vs. 50 mM H_2_O_2_. Different colors for fonts identify rescaled GO terms with fold change ≥ 1 at p < 0.05, determined by t-test: red, significantly expressed in both; blue, significantly expressed in treatment mentioned on the x-axis; cyan-green, significantly expressed in treatment mentioned on the y-axis; and black, no significant expression in either. (Note: * represents scaling by dividing the abundances for the treatment with the corresponding values for HDO_(MS)_).

The metaproteomics analysis of FS bioreactors revealed protein expressions related to metabolism (nitrogen metabolism, oxidative phosphorylation), genetic information processing, and environmental information processing (Figure S5). Functional analysis demonstrated significant upregulation of oxidative stress responses in 5×10^-5^ M L-NE cultures compared to dextrose, particularly catalase-peroxidase and thioredoxin disulfide reductase activities (Figure 2). The enhanced glutamine-ammonia ligase activity in L-NE cultures indicates altered nitrogen metabolism through flux redirection to glutamate/glutamine synthesis. Thioredoxin reductases maintain enzyme catalytic activity by repairing oxidative stress-induced disulfide bonds (Arts et al., 2016), while thioredoxin peroxidase employs thioredoxin to detoxify H_2_O_2_ (Somprasong et al., 2012). Notably, 50 µM H_2_O_2_ treatment induced higher superoxide dismutase expression than 5×10^-5^ M L-NE, catalyzing superoxide (O_2_ ^·-^) conversion to O and H O. Enhanced catalase-peroxidase and thioredoxin reductase activities in L-NE versus H_2_O_2_ cultures suggest *oxyR* activation from increased H_2_O_2_ production, consistent with our ROS gene abundance findings (Figure S4). L- NE and dextrose treatments showed distinct proteomic profiles despite similar community compositions. The significant upregulation of antioxidative enzymes (catalase, thioredoxin, superoxide dismutase) in L-NE-treated cultures indicates sufficient ROS stress to activate *soxRS* and *oxyR* genes and their corresponding proteins (Figure S4). L-NE treatment also enhanced nitrogen reductase and glutamate-ammonia ligase expression, suggesting potential applications in community nitrogen metabolism for resource recovery. The increased expression of oxidoreductases, monooxygenases, copper ion binding laccases, and heme binding cytochromes in L-NE versus dextrose treatments indicates potential roles for *Pseudomonadaceae* in organic micropollutant removal. These findings demonstrate that while L-NE and dextrose support similar community compositions, they induce distinct enzymatic responses, with L-NE eliciting more sophisticated ROS management strategies than either dextrose or H_2_O_2_ treatments.

### 3.4 Mechanistic differences between L-NE and dextrose treatments by M-model prediction of metabolic activities

Given the dominance of *Pseudomonadaceae* under both L-NE and dextrose conditions (Figure S3) and our observed differences in oxidative stress responses and protein expression, we employed metabolic modeling to elucidate the underlying mechanistic distinctions. Using *Pseudomonas putida* KT2440 as a model organism (Nogales et al., 2020), we performed flux balance analysis (FBA) with a proteome-constrained modified genome-scale metabolic model (M-model). A custom metaproteome-based flux scaling pipeline was developed, incorporating protein constraints and archetypal analyses to identify key metabolic differences between treatments. M-model simulations revealed distinct regulation patterns in several central metabolic pathways, including gluconeogenesis/glycolysis, oxidative phosphorylation, TCA cycle, sulfur metabolism, and glutamate metabolism between L-NE and dextrose-treated cultures (Figure 3, Supplementary Data 3). L-NE cultures exhibited significantly higher oxygen consumption compared to dextrose treatments, attributed to enhanced terminal oxidase activity and subsequent ATP production (Figure 4 and Figure S6). This observation aligns with previous findings that lower TCA cycle intermediates enhance oxygen consumption and proton motive force in stationary phase cells (Meylan et al., 2017). The model also supports established evidence that *Pseudomonas* demonstrates higher growth rates when supplied with TCA intermediates like succinate versus dextrose (Nikel et al., 2015, 2014; Tiwari and Campbell, 1969). Moreover, L-NE degradation has been reported for *Pseudomonas* species (Cuskey et al., 1987; Cuskey and Olsen, 1988; Luengo and Olivera, 2020). While L-NE may not serve as an energetically optimal carbon source, its potential role in providing TCA cycle intermediates could explain the enhanced microbial growth observed in complex media. We acknowledge that additional factors, such as autoinducers, might contribute to L-NE-enhanced growth rates, though these aspects exceed current modeling capabilities.

**Figure 3.**
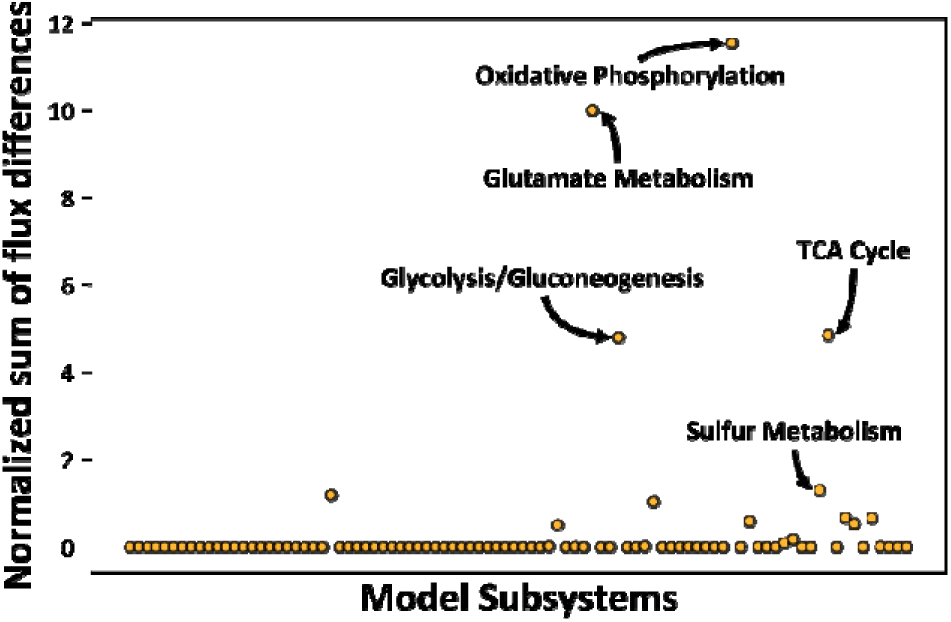
A scatterplot of the predicted normalized fluxes. To determine the subsystems that were differentially changed between dextrose and L-NE conditions, the absolute flux values were normalized, and the difference was computed between L-NE and dextrose. Following this, the absolute difference was summed up for each subsystem. The top five subsystems that were altered between the two conditions are shown in the figure. Notably, the L-NE degradation pathway was removed from the subsystem analysis.

**Figure 4.**
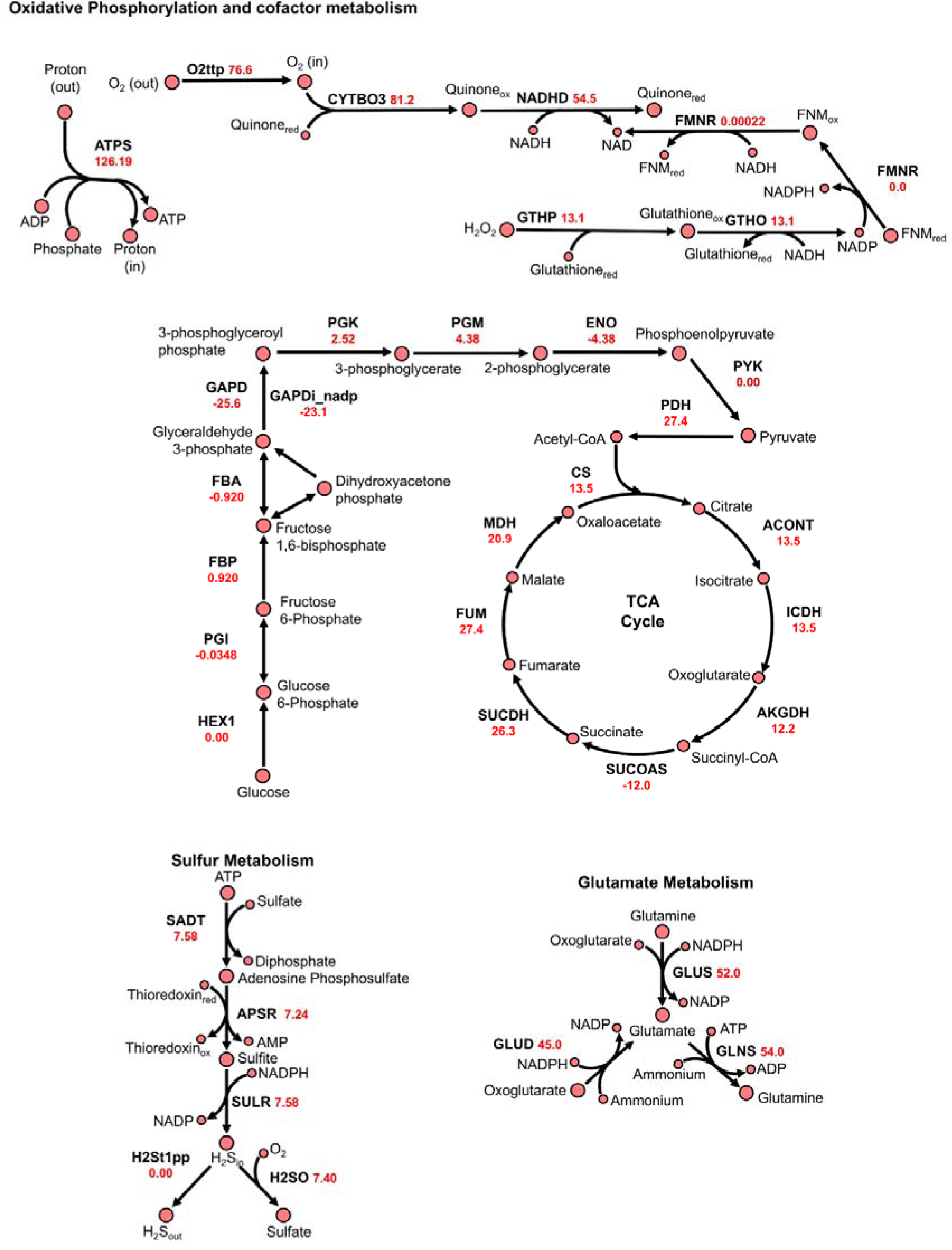
Pathway maps showing normalized fluxes for L-NE treatments. The higher growth rate in the L-NE treatment compared to dextrose (Figure S6) results from increased O_2_ consumption, which shifted the fluxes towards greater terminal oxidase and ATP synthase activity. L-NE additionally reprograms the cofactor metabolism, TCA cycle, Glycolysis/Gluconeogenesis, sulfur metabolism, and thioredoxin-based and glutathione-based oxidative stress mitigation strategies. In the model, reaction GLNS and FMN reductase reactions are split into two; the pathway map shows only one of these fluxes.

Consistent with our experimental findings in previous sections, the model predicted stronger oxidative stress mitigation responses under L-NE treatment compared to dextrose, particularly through elevated glutathione-based peroxidase activity. The predicted alterations in sulfur metabolism align with previously documented responses to H_2_O_2_ and PQ-induced oxidative stress in *Pseudomonas* strains (Bojanovič et al., 2017; Hare et al., 2011; Palma et al., 2004; Yeom et al., 2012). The enhanced utilization of redox metabolites (thioredoxin and glutathione) correlates with increased sulfur metabolism under L-NE conditions, supporting our experimental observations of enhanced ROS management.

The model further validated our metaproteomic findings regarding nitrogen metabolism, showing L-NE-specific effects through the glutamate/glutamine pathway. Pathway maps revealed L-NE-induced upregulation of the TCA cycle and associated oxidative phosphorylation pathways (Figure 4, Figure S6 and Supplementary Data 4). The increased flux through these pathways, particularly terminal oxidase reactions, explains the elevated oxygen consumption and ATP synthesis rates. The L-NE degradation pathway’s connection to NADH cofactor regeneration influences ubiquinone and NADP cofactor cycling, providing a mechanistic basis for the observed metabolic shifts.

Supporting our metaproteomic data, the simulations confirmed L-NE’s distinct effects on nitrogen metabolism (glutamate/glutamine synthesis) and carbon-associated (TCA cycle) pathways compared to dextrose. The model particularly highlighted *Pseudomonas*’s preferential use of glutathione-based mechanisms for oxidative stress elimination, linking this to the observed sulfur metabolism reprogramming (Figure 3). These insights into L-NE-induced metabolic reprogramming suggest potential applications for enhancing resource recovery in microbial communities.

### 3.5 L-NE Stimulates Quorum Sensing and Virulence Traits

Our preceding analyses demonstrated distinct metabolic and stress response mechanisms between L-NE and dextrose treatments despite similar community compositions. To investigate the potential virulence implications of these differences, we performed growth assays examining autoinducer effects in conditioned media generated at 8 and 24 hours through pour plate analysis following 24 hours of incubation. The serial dilution of L-NE and dextrose-conditioned medium demonstrated maximum autoinducer growth activity at 0.6% (v/v) conditioned media. The conditioned media from L-NE exposed mixed cultures showed substantially higher bacterial growth (10^6^-10^7^ CFU ml^-1^) at both 8 and 24 hours compared to dextrose cultures (10^4^ CFU ml^-1^). To establish that enhanced bacterial growth resulted from autoinducer production, we determined *qseC* and *luxS* expression at 24 hours using real-time quantitative PCR (qPCR). The relative abundance of both *qseC* and *luxS* was higher in L-NE samples compared to dextrose-spiked cultures, indicating that L-NE might activate LuxS expression to synthesize autoinducers (AI-3) during the exponential growth phase, suggesting cell-to-cell signaling as a mechanism for enhanced bacterial growth (Figure S7). The absence of elevated autoinducer genes in dextrose treatments revealed its primary role as a carbon source, while L-NE promoted growth through both metabolic utilization and autoinducer triggering. This mechanistic distinction is particularly significant when considered alongside our previous observations of enhanced oxidative stress tolerance, distinct proteomic profiles indicating upregulation of both stress response and metabolic pathways, and species-level metabolic reprogramming that may facilitate both enhanced growth and increased signaling molecule synthesis. While L-NE exposure did not preferentially enrich known pathogenic taxa in our experimental communities, the enhanced quorum sensing activity coupled with altered stress responses and metabolic adaptations suggests the potential for increased virulence traits in wastewater communities.

## 4. Conclusion

This study provides the first comprehensive characterization of L-NE’s effects on complex wastewater microbial communities, revealing sophisticated metabolic and regulatory mechanisms that extend beyond previously documented pure culture responses. Our multi-tiered experimental approach, integrating taxonomic profiling, targeted metaproteomics, and metabolic modeling, uncovered three key findings: First, we demonstrated L-NE’s unique dual role as both a metabolic substrate and signaling molecule in mixed microbial consortia. Despite containing 10-fold less carbon, L-NE treatments achieved higher growth (10L CFU mLL¹) than dextrose while maintaining similar community compositions dominated by *Pseudomonadaceae*. This enhanced growth was mechanistically linked to L-NE’s ability to provide TCA cycle intermediates and stimulate oxidative phosphorylation, as revealed by our proteome-constrained metabolic modeling. Second, we identified a sophisticated oxidative stress response network in L-NE-exposed communities. The coordinated upregulation of oxidative stress genes (*oxyR*, *soxRS*) and antioxidant enzymes, particularly through enhanced glutathione-based mechanisms, demonstrates community-level adaptation to L-NE-induced ROS. Notably, this stress response was distinct from both dextrose and HLOL controls, indicating L-NE-specific regulatory pathways. Third, while L-NE did not enrich pathogenic taxa, it significantly increased autoinducer gene (*luxS*, *qseC*) abundance and enhanced quorum sensing activity compared to dextrose controls. This observation, coupled with distinct metabolic reprogramming and stress responses, suggests the potential for increased virulence traits without requiring changes in community composition. Our findings highlight the need to consider neuroendocrine compounds in wastewater treatment design - compounds that enter wastewater through diverse sources, including plants and fecal waste, with concentrations potentially elevated during periods of infection and stress - particularly their impacts on community function and virulence trait expression. These integrated findings establish L-NE as a significant modulator of microbial community dynamics in engineered ecosystems. Future research should focus on long-term adaptation studies, examination of horizontal gene transfer rates under enhanced quorum sensing conditions, and detailed assessment of virulence factor expression in specific taxa under L-NE exposure. The mechanisms elucidated here provide new insights for understanding host-microbe interactions in environmental systems and may inform strategies for optimizing wastewater treatment processes to address emerging contaminants.

## Acknowledgments

This project was partially funded by an award from the Royal Society of New Zealand’s Marsden Fund (Contract Number: MFP-UOA2018). The authors thank the Gavin Lear group for performing the microbial taxonomic analyses.

## CRediT authorship contribution statement

**Amrita Bains:** Writing – original draft, Methodology, Investigation, Visualization, Formal analysis. **Sanjeev Dahal:** Writing – original draft, Methodology, Visualization, Formal analysis. **Bharat Manna:** Writing – original draft, Methodology, Visualization, Formal analysis. **Mark Lyte:** Writing – review & editing, Conceptualization. **Laurence Yang:** Writing – review & editing, **Naresh Singhal:** Writing – review & editing, Supervision, Funding acquisition, Conceptualization.

