## Supplementary Information for "L-norepinephrine Induces Community Shift, Oxidative Stress Response, Metabolic Reprogramming, and Virulence Potential in Wastewater Microbiomes"

### for

#### S1. Experimental Methodology

##### S1.1 Reactor operational parameters and sampling

The mixed liquor sludge from a dairy farm was washed three times by mixing and decanting 0.5 L of supernatant with distilled water (Milli-Q system, Millipore, Darmstadt, Germany) in volumetric cylinders to achieve volatile suspended solids (VSS) of 3 g-VSS/L. The synthetic wastewater feed consisted of sodium acetate trihydrate (63 mM), magnesium sulfate heptahydrate (3.6 mM), potassium chloride (4.7 mM), ammonium chloride (35.4 mM), di-potassium hydrogen phosphate (4.2 mM), potassium dihydrogen phosphate (2.1 mM) and 10 mL/L of a trace element solution [1], all purchased from Sigma Aldrich; purity ≥ 98% (St. Louis, MO, USA). Oxygen (99%) was purchased from BOC (Auckland, New Zealand). The total COD in each reactor is 500 mg-COD/L of which 400 mg-COD/L comes from acetate and 100 mg-COD/L from methanol (Bassin et al., 2011). A composition of 500 mg-COD/L organic carbon and 60 mg-N/L ammonia is representative of wastewater in activated sludge systems, although it is at the high end of the concentration range.

##### S1.2. Real-time quantitative PCR

The abundance levels of four target genes (*soxRS, oxyR*, *luxS* and *qseC*) were analyzed with a qPCR sequence detection system (QuantStudio 5 Real-Time PCR System). Total bacterial genomic DNA was extracted from sludge samples (1 mL) using a PowerSoil DNA isolation kit (MoBio, Carlsbad, USA) following the manufacturer’s protocol. The housekeeping gene (16S rRNA) gene was used as a reference for the quantification of the genes expressed during endogenously expressed stress. qPCRs were done in a final volume of 10 µl containing: 5 µl Power SYBR® Green PCR master mix, 0.5 µl each of 10 µM forward and reverse primers (listed in Table S5), and 2 µl of extracted genomic sample DNA. Gene amplification was analyzed by measuring fluorescence. Reaction mixtures were prepared in a 384-well plate (Raylab NZ Ltd. Cat# CP3436.S) using a single channel multi-pipette (for master mix without DNA samples). The plate is placed in a 384-well plate base and sat on ice the whole time. After adding the master mix and DNA samples to the plate wells, the plate was sealed with a MicroAmp™ Optical Adhesive Film (Applied Biosystems, Cat#4311971) with an applicator. The plate was then centrifuged (Sigma 2.5 centrifuge) for 2-3 min at max speed to mix samples and remove bubbles. The real-time PCR was carried out using a 7900HT sequence detection system with SDS 2.3 software (Applied Biosystems). A real-time PCR machine was connected to the computer using a cloud account, and the run was set up. The experimental type and the chemistry were changed to comparative CT and SYBR green. Sample wells were labeled, and the default thermal cycler program was used: (i) 50˚C for 2 min, 95˚C for 10 min, (ii) 40x95˚C for 15 sec and 60˚C for 2 min, (iii) 95˚C for 15 sec, 60˚Cfor 15 sec and 95˚C for 15 sec). The program was saved, and the plate was inserted into the PCR machine to run.

**S1.3. Protein extraction and identification**

Proteomic analyses were performed to screen for oxidative stress-associated enzymes as well as non-ROS enzymes. Protein extraction was performed by cell lysis of farm and municipal sludge cultures. 1 ml of sludge sample was pelleted and washed twice with 50 mM ammonium bicarbonate and spun down at 16,000 x g for 5 min at 4°C. Pellets were resuspended in 150 µl 7M urea-thiourea buffer and sonicated for 15 s (three rounds on ice). After sonication, 500 µL of 15% chilled trichloroacetic acid (TCA) was added, vortexed well, and incubated for 15 min. at 4˚C. The samples were then centrifuged at 16,000 x g for 15 min and the supernatant containing TCA was removed. Pellets were washed with 1 mL acetone and centrifuged at 16,000 x g for 5 min and dried completely. The dried pellet was again resuspended in 150 µL 7M urea-thiourea buffer and sonicated for 15 s (three rounds on ice). The sonicated samples were reduced at 56˚C for 20 min. on heat block and pH was checked after reduction. The reduced samples were then alkylated by adding 50 mM iodoacetamide (IAM) and incubated in dark for 30 min. The reactions were quenched by adding 10 mM dithiothreitol (DTT) and samples were centrifuged at 16,000 x g for 5 min and the supernatant was collected. The protein concentration was quantified by performing an EZQ protein assay using an EZQ protein quantification kit following the manufacturer's protocol. The protein concentrations were normalized to 30 µg/µL. The protein digestion was performed by adding trypsin (1 µg/µL) for 2 hours at 45˚C in the microwave. The acidification of digested samples was done with 50% formic acid and pH was checked (2-3). Samples were centrifuged at 16,000 x g for 3 min. The proteins were extracted by solid-phase extraction. 10 mg Oasis HLB cartridges were pre-conditioned with 0.5 mL each 100% methanol and 0.1% formic acid. The sample supernatant was loaded on the pre-conditioned cartridges. After sample elution, cartridges were washed with 1 mL of 0.1% formic acid. The proteins were eluted in 1.5 mL Eppendorf tubes with 300 µL of 50% acetonitrile in 0.1% formic acid. The eluted proteins were speed-vacuumed to 15-20 µL as the final volume. A 10 ul aliquot of each sample was injected onto a 0.3x 10mm trap column packed with Reprosil C18 media (Dr Maisch) and desalted for 5 minutes at 7ul/min before being separated on a 0.075 x 200 mm picofrit column (New Objective) packed in-house with Reprosil C18 media. The following gradient was applied at 300nl/min using a NanoLC 400 UPLC system (Eksigent): 0min 1%B; 2min 1%B; 105min, 35%B; 110min, 98%B;

115min, 98%B; 116min, 1%B; 125min, 1%B, where A was 0.1% formic acid in water and B was 0.1% formic acid in acetonitrile. The picofrit column spray was directed into a TripleTOF 6600 Quadrupole-Time-of-Flight mass spectrometer (Sciex) scanning from 350-1600 m/z for 250ms, followed by 45ms MS/MS scans on the 45 most abundant multiply-charged peptides (m/z 100-1600) for a total cycle time of ~2 seconds. The mass spectrometer and UPLC system were under the control of the Analyst TF 1.7 software package (Sciex). The resulting data from each band were searched against a database containing the Uniref Bacteria sequences with 50% identity (Uniprot.org) using Protein Pilot version 5.0 (Sciex). Search parameters were as follows: Sample Type, Identification; Search Effort, Thorough; Cys Alkylation, Iodoacetamide; Digestion, Trypsin; ID Focus, Biological Modifications, Amino Acid Substitutions. Peptide and Protein summary files were exported for further analysis.

##### S1.4. Metaproteome Data Analysis

Two separate pipelines for matching peptides were pursued to identify functions related to specific peptides. In the first pipeline, all 40 detected peptides were searched against the Uniprot database using the Peptide Search tool (https://www.uniprot.org/peptidesearch/). The data from Peptide Search was filtered for the 127 taxonomical families that were detected in the metagenome analysis. That is, any function associated with proteins from non-detected families was excluded. In this process, two peptides (GATVMISPYVMHR, WSEQGAAPASHLR) were excluded as their associated functions mapped to family not identified in the metagenomic data. Under the assumption that peptide detection corresponds to functional activity, these peptides and their respective functions were manually added back for the subsequent analyses.

In the second pipeline, the peptides were assigned to gene ontology (GO) terms using Unipept [2] through the proteomics Analysis tool (https://unipept.ugent.be/datasets). Default parameters were applied except the equating isoleucine and leucine option was unchecked. The Unipept results (molecular function, cellular component, and biological process) for all the peptides were then merged into one table. Any peptide sequence modified by the Unipept to search its database was reverted back to its original sequence in the final results to seamlessly intersect the results of both pipelines (Supplementary Data 1). Using the intersection table, the molecular functions (GO terms) of the microbial communities were analyzed under different treatment conditions. First, the peptide intensities for the mapped GO terms were summed and the average of the logs of the total intensities for biological replicates was calculated. After the data normalization with the HDO condition, a two-sided T-test was performed to identify the functions that changed significantly between different conditions. For the T-test, the null hypothesis that the averages of two independent samples are identical was assumed. To visualize the results of the analyses, scatterplots were utilized.

**S1.5. M-model simulations of archetypes, analysis and visualisation**

Archetypal analysis (AA) finds "archetypes" which are the points within the multivariate data set whose convex combination can well represent the data set [3]. It can be viewed as a dimension-reduction method such that each archetype refers to a distinct state of cell physiology. In this analysis, a dataset X can be approximated as

$$X \approx ZA,$$

where Z is the matrix of archetypes and A is a matrix of coefficients such that $A_{ij} \geq 0$ and

$\Sigma_{pj=1} \Sigma j=1p$

$A_{ij} = 1$ for *p* archetypes.

Likewise, Z can be constrained as

$Z = XB$ where B is the coefficient matrix such that $B_{ij} \geq 0$ and

$\Sigma_{pj=1} \Sigma j=1p$

$B_{ij} = 1$ for *p* archetypes such that Z is also constrained to be a convex combination of the data points in X.

The archetypal analysis was performed using *py_pcha* tool, which is a Python implementation of principal convex hull analysis (https://github.com/ulfaslak/py_pcha). The matrix used for the analyses was derived by performing simulations on a model of *Pseudomonas putida* KT2440 (*i*JN1463) constrained using targeted proteome data [4]. This model was used to study L-NE utilization in *Pseudomonas* species, which were the most abundant members of the microbiome.

We made two assumptions guided by the literature to modify this model. First, *P. putida* KT2440 is unable to catabolize several biogenic amines [5], but L-NE degradation has been reported for several *Pseudomonas* species, so we added putative reactions for L-NE transport and metabolism for a related *Pseudomonas* strain, *Pseudomonas putida* U, to the model [6–8]. Second, as *Pseudomonas putida* cannot utilize methanol as the sole carbon source for growth (Hintermayer and Weuster-Botz 2017), we treated the reaction catalyzed by formaldehyde dismutase as irreversible to prevent methanol use for growth, consistent with the MetaCyc database [9].

We performed model simulations to optimize the growth rate under the following fluxes (shown in mmol $gDW^{-1} hr^{-1}$) for the following substrates: 10 for acetate, 18 for oxygen, 100 for ammonium, and 1000 for methanol, sodium, phosphate, chloride, magnesium, calcium, potassium, iron, copper, sulfate, manganese, molybdenum, zinc, cobalt, nickel, water, and proton. Most fluxes were chosen based on the previous studies on M-models and on their concentration in the medium [10]. The oxygen uptake rate constraint was approximated using another study [11]. The fluxes for acetate, ammonium, and methanol were arbitrarily chosen based on their concentration in the experimental medium. The uptake fluxes for L-NE and dextrose for their respective medium were constrained at 6 mmol $gDW^{-1}$ $hr^{-1}$, which was the default uptake rate for dextrose set in the model.

Then, we performed a parsimonious flux balance analysis (pFBA) [12] using this medium for the HDO condition, i.e., without dextrose or L-NE in the medium. The proteins from the targeted proteome data were retrieved for *Pseudomonas putida* (Supplementary Data 2), and the reactions catalyzed by those proteins were identified. Following this, the $k_{effs}$ for each reaction-gene pair (associated with these proteins) were computed using the formula:

$$k_{eff} = (v_{pfba}/i) * N$$

where $v_{pfba}$ is the flux computed by pFBA for the HDO condition, $i$ is the protein intensity, and $N$ is the number of reactions catalyzed by a particular gene (Supplementary Data 2) for details on the proteins and reactions constrained). Computed values of zero for $k_{eff}$ were replaced with the lowest computed non-zero value. Subsequently, a target flux analysis was performed using HDO with dextrose or L-NE added at 6 mmol $gDW^{-1}$ $hr^{-1}$ for the reaction-gene pairs (specified above) using the formula:

$$v_t = k_{eff} * (i / N)$$

To obtain the target flux ($v_t$), which is influenced by peptide intensity ($i$) for certain growth conditions, for a specific reaction, the gene-reaction rules (e.g., linking gene *a* with gene *b* using AND or OR) were parsed such that for AND take the lower of the two genes and for OR take the sum of the fluxes associated with these genes. Following the flux calculation, we performed a minimization of the sum of fluxes for these reactions (associated with proteins identified in proteomics) at fixed growth rates ($\mu$) ranging from zero to the optimal growth rate (identified by FBA), i.e., 100 simulations, using the following formula:

$$z = \sum(v - v_t)^2$$

where *z*, the sum of the square of errors, is the objective function, and *v* is the flux obtained by simulating a particular condition. We then performed another pFBA analysis at different growth rates (zero to FBA optimal growth rate, 100 simulations) by bounding the fluxes of the reactions catalyzed by the proteins identified in the proteome data using the fluxes predicted in the above minimization procedure. These simulations provided the matrix containing rows (growth rates) and columns (reaction fluxes) that was then used in the subsequent archetypal analysis to identify representative archetypes of flux distribution [3].

We chose two archetypes for the AA using the elbow method on the scree plot. These archetypes could explain >96% of the variance in both conditions (Figure S8, Supplementary Data 5). There were two other criteria for choosing the number of archetypes: 1) The growth rate in L-NE is higher than that in dextrose, and 2) the sum of squared error should be minimum as indicated by the elbow of the scree plot (Figure S9). In AA restricted by these points, we chose one archetype each at the highest growth rates of ~0.060 $hr^{-1}$ and ~0.083 $hr^{-1}$ for dextrose and L-NE growth conditions, respectively. These two archetypes (one for each condition) reflect the best approximation of the growth and proteomic assays. We next compare these two archetypes in the subsequent analyses and visualization.

We used the Escher package (v. 1.7.3) in Python (v. 3.7.7) to visualize the maps of reaction fluxes normalized with the corresponding biomass flux [13]. Normalization was performed to compare the proteome-constrained fluxes independent of the differences in growth rates between different conditions. For visualizing pathways, we modified the maps created by the Escher online tool using Affinity Designer (v. 1.9.3) and Inkscape (v. 1.1.2). Genome-scale models typically contain information about subsystems; this knowledge was used to predict which subsystems were altered between two conditions. For the added pathways of L-NE transport and degradation, we assigned them the subsystem 'L-NE_Degradation' within the modified model (see 'M-model simulations of archetypes' section). Two modifications were made to the subsystems: 1) LNE_Degradation subsystem was removed as the pathway was assumed to be active in only L-NE growth condition, and 2) Glycolysis and Gluconeogenesis were assigned to a new subsystem Glycolysis/Gluconeogenesis to avoid any confusion on the direction of the fluxes. Following these changes, we identified subsystems showing the most change by dividing the sum of the absolute difference in the normalized fluxes for two conditions by the number of reactions in the subsystem.

A few assumptions had to be made to use the P. putida model to study the global effects of catecholamines (L-NE) versus dextrose. *P. putida* KT2440 naturally does not possess the machinery to catabolize several biogenic amines [5]. The core *Pseudomonas putida* model seems to contain the same gene content as in at least 95% of other *Pseudomonas putida* reconstructions[4]. With this consideration, and because the computational analyses were performed assuming the genome-scale model is representative of the community *Pseudomonas strains,* the L-NE degradation pathway of the related *Pseudomonas* strain, Pseudomonas putida U was added to the model in order to study the effect of L-NE on the Pseudomonas strains of the microbial community. Another consideration while constraining the model was that the proteomic data were available for 24-hour cultures, at which point the growth rates were very low (~0.01-0.04). To make sure the proteome-constrained fluxes were comparable between L-NE and dextrose cultures, the fluxes were normalized by the respective growth rates. Further, to remove the effect of methanol utilization and to focus on the effect of L-NE and dextrose, the reaction catalyzed by formaldehyde dehydrogenase was made irreversible.

**Table S1.** Description of experimental conditions evaluated in this study.

| Experimental Condition | Description |
|---|---|
| High Dissolved Oxygen (HDO) | Base bioreactor with dairy farm and activated sludge cultures maintained in synthetic wastewater under constant aeration at ~8 mg/L of dissolved oxygen. |
| $H_2O_2$ | The activated sludge culture maintained in synthetic wastewater media containing 5, 10 and 50 μM hydrogen peroxide as ROS control. |
| Dextrose | HDO cultures supplemented with $1x10^{-5}$, $5x10^{-5}$, and $1x10^{-4}$ M dextrose to serve as metabolic carbon source control. |
| L-NE | HDO cultures supplemented with $1\times10^{-5}$, $5\times10^{-5}$ and $1\times10^{-4}$ M L-NE. |

**Table S2.** Details of sampling number and intervals.

| Experiments | Number of samples | | Sampling intervals (hours) |
|---|---|---|---|
| | Biological Replicate | Technical Replicate | |
| Bacterial growth | 2 | 2 | 0, 8 and 24 |
| Gene expression | 2 | 2 | 24 |
| Proteomics | 2 | 2 | 24 |
| Microbial speciation | 2 | 2 | 24 |

**Table S3.** Mixed culture growth curve (CFU/mL) under different Dextrose and L-NE concentrations.

| Time (Hr) | Farm Sludge (Live Biomass) | | | | | | Municipal Sludge (Live Biomass) | | | | | |
|---|---|---|---|---|---|---|---|---|---|---|---|---|
| | Dextrose | | | L-NE | | | Dextrose | | | L-NE | | |
| | $1\times10^{-5}$ M | $5\times10^{-5}$ M | $1\times10^{-4}$ M | $1\times10^{-5}$ M | $5\times10^{-5}$ M | $1\times10^{-4}$ M | $1\times10^{-5}$ M | $5\times10^{-5}$ M | $1\times10^{-4}$ M | $1\times10^{-5}$ M | $5\times10^{-5}$ M | $1\times10^{-4}$ M |
| **0** | $1x10^2$ | $1x10^2$ | $1x10^2$ | $1x10^2$ | $1x10^2$ | $1x10^2$ | $1x10^2$ | $1x10^2$ | $1x10^2$ | $1x10^2$ | $1x10^2$ | $1x10^2$ |
| **8** | $3x10^4 \pm 3x10^3$ | $6x10^3 \pm 2x10^2$ | $2x10^4 \pm 2x10^3$ | $6x10^5 \pm 2x10^4$ | $5x10^6 \pm 2x10^5$ | $3x10^6 \pm 2x10^5$ | $6x10^4 \pm 7x10^3$ | $5x10^4 \pm 7x10^3$ | $6x10^5 \pm 7x10^3$ | $6x10^4 \pm 2x10^3$ | $3x10^5 \pm 1x10^4$ | $6x10^4 \pm 2x10^3$ |
| **24** | $3x10^5 \pm 6x10^3$ | $3x10^5 \pm 7x10^3$ | $3x10^5 \pm 4x10^3$ | $2x10^6 \pm 2x10^4$ | $1x10^8 \pm 4x10^6$ | $2x10^7 \pm 3x10^5$ | $3x10^6 \pm 2x10^4$ | $1x10^5 \pm 5x10^3$ | $3x10^6 \pm 4x10^4$ | $4x10^6 \pm 5x10^5$ | $2x10^7 \pm 5x10^5$ | $8x10^6 \pm 3x10^5$ |

**Table S4.** Standard Error of the slopes as shown in Figure 1.

| | Equation | Slope | Standard Error |
|---|---|---|---|
| **a** | $Y$=1.07$x$+0.15 | 1.07 | 0.05 |
| | $Y$=1.05$x$+0.07 | 1.05 | 0.055 |
| | $Y$=0.96$x$+0.33 | 0.96 | 0.053 |
| **b** | $Y$=0.65$x$+0.49 | 0.65 | 0.058 |
| | $Y$=0.83$x$ −1.38 | 0.83 | 0.134 |
| | $Y$=0.78$x$ −1.41 | 0.78 | 0.21 |
| **c** | $Y$=0.003$x$ −0.23 | 0.003 | 0.05 |
| | $Y$=0.03$x$−0.12 | 0.03 | 0.048 |
| | $Y$=0.25$x$+0.28 | 0.25 | 0.094 |

**Table S5.** Primer sequences of genes activated in oxidative stress and the genes activating the production of autoinducers by bacterial species.

| | | | | | |
|---|---|---|---|---|---|
| **Bacteria** | All Bacteria | 16S rRNA | 16S rRNA-F<br>16S rRNA-R | 5'-TCGTCGGCAGCGTCAGATGTGTATAAGAGACAGCCTACGGGNGGCWGCAG-3'<br>5'-GTCTCGTGGGCTCGGAGATGTGTATAAGAGACAGGACTACHVGGGTATCTAATCC-3' | Quast et al., 2013 |
| | All Bacteria | *oxyR* | UR029-F<br>UR084-R | 5'- CCGGAATTCTCTGGCGTCAATTAT-3'<br>5'- ACCACCTTTAACTACCCGACG-3' | Daugherty, Suvarnapunya, & Runyen-Janecky, 2012 |
| | All Bacteria | *soxRS* | UR027-F<br>UR028-R | 5'- CCGGAATTCTGAGCATGCATTTCTTG-3'<br>5'- CGCGGATCCGCTTTAGTTTTGTGTTTGC-3' | Daugherty et al., 2012 |
| | *Vibrio harveyi* | *luxS* | LuxSFHis<br>LuxSRHis | 5'-GGTACCCCGTTGTTAGATAGCTTCAC -3'<br>5'-AAGCTTCTAGATGTGCAGTTCCTGCAACT -3' | Sperandio et al., 2003 |
| | *Enterohemorrhagic E. coli* | *qseC* | YfhKFBD<br>YfhKRBD | 5'-GGTACCTATCTGAACTTCCCCTCGGTT -3'<br>5'-GAATTCCCTTTCGTGTTTTTCGACGACGG -3' | Reading et al., 2007 |

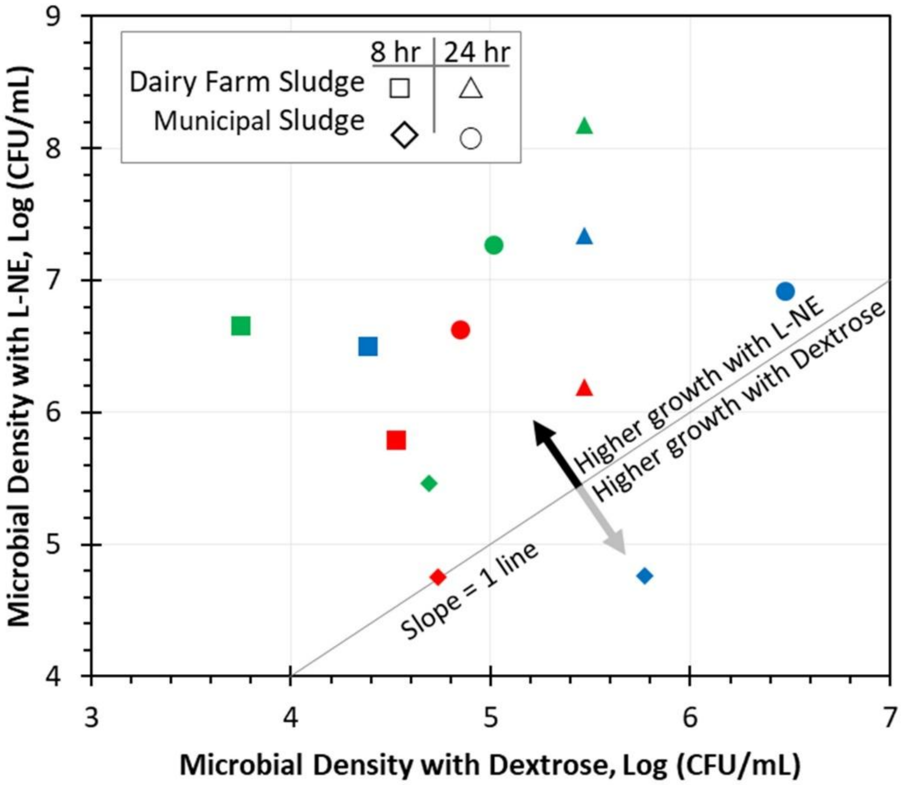


**Figure S1.** Microbial growth with different concentrations of L-NE ($1x10^{-5}$ M, $5x10^{-5}$ M, and $1x10^{-4}$ M), and Dextrose ($1x10^{-5}$ M, $5x10^{-5}$ M, and $1x10^{-4}$ M) over 8 hr and 24 hr in MS and FS cultures. The red, green, and blue colors represent $1x10^{-5}$ M, $5x10^{-5}$ M, and $1x10^{-4}$ M concentrations, respectively.

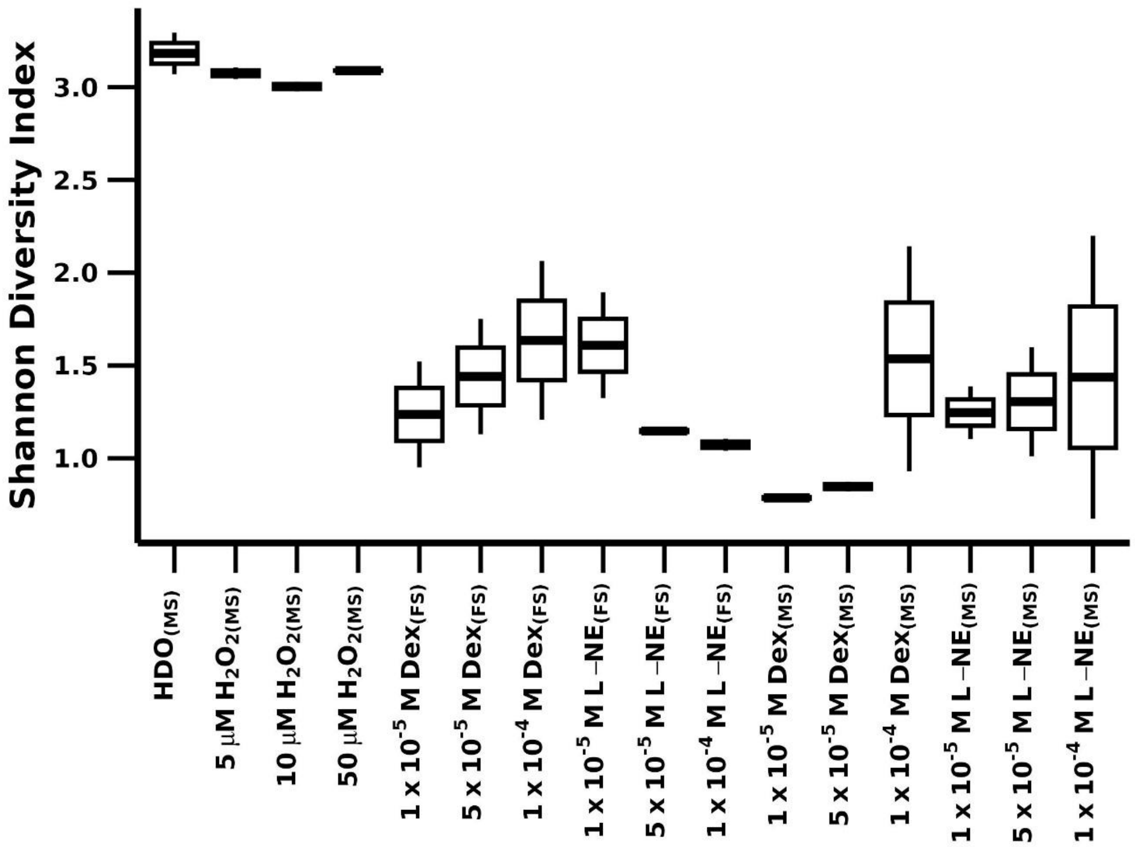


**Figure S2.** Shannon diversity of the microbial population in different treatment systems with HDO, $H_2O_2$, dextrose, and L-NE.

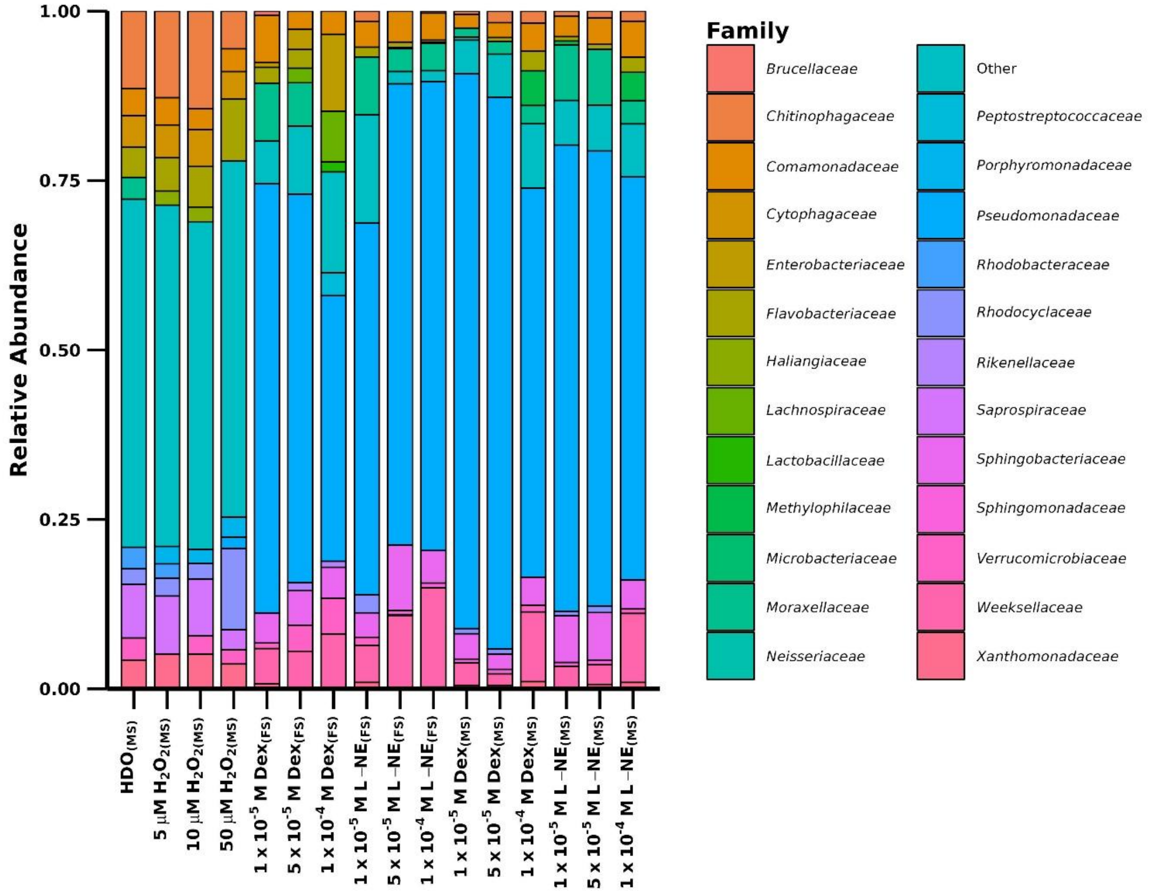


**Figure S3.** 16S rRNA-based relative abundance of the microbial families in different treatment systems with HDO, $H_2O_2$, dextrose, and L-NE.

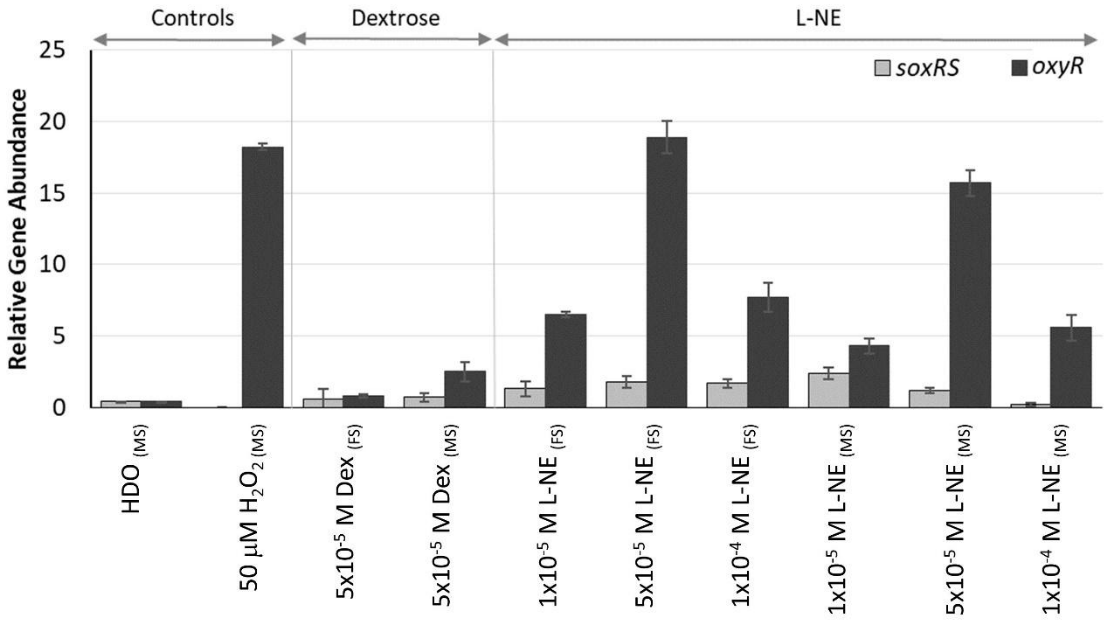


**Figure S4.** Relative abundances of *soxRS* and *oxyR* genes, relative to 16S genes, in HDO, 50 μM $H_2O_2$ in municipal sludge, and dextrose and L-NE in both farm and municipal sludge cultures (n = 4; $p < 0.05$; mean ± standard deviation).

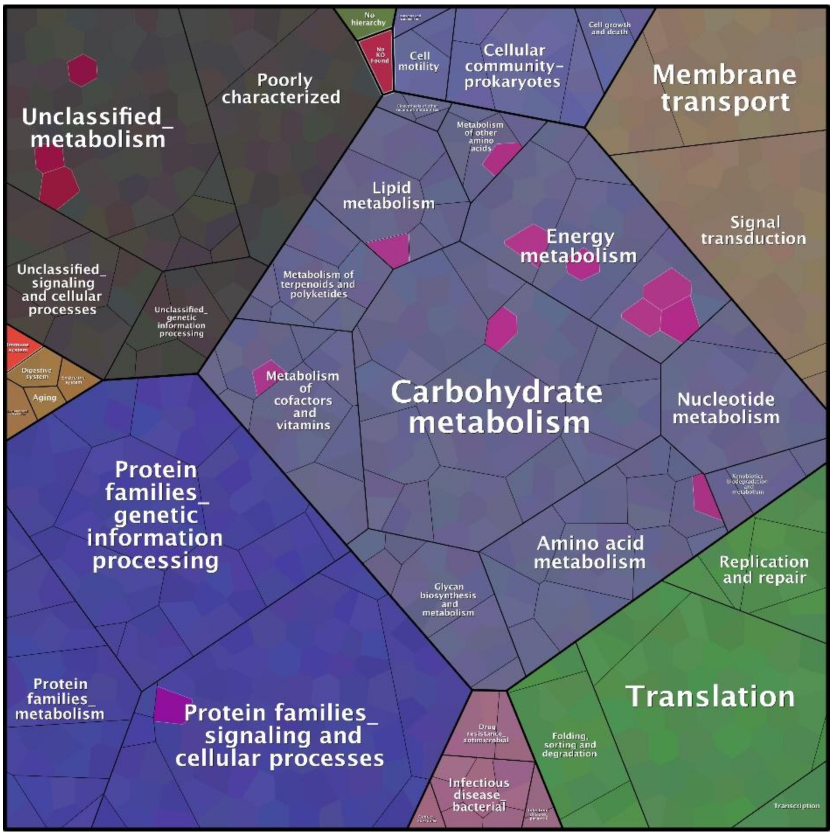


**Figure S5.** Proteomaps visualizing the pathways detected in the global proteomic analysis. A functional overview illustration of the community proteome. The black sectors show functions that could not be categorized into distinct KEGG hierarchies and are binned into generic functions. The brown sector shows proteins with absent KEGG orthology (KO) that are grouped as "No KO Found". The dark green sector shows functions with KO but absent hierarchies in the template treemap and grouped as "No hierarchy". The dark pink sectors with white borders represent stress-related functions characterized using targeted proteomics in this study.

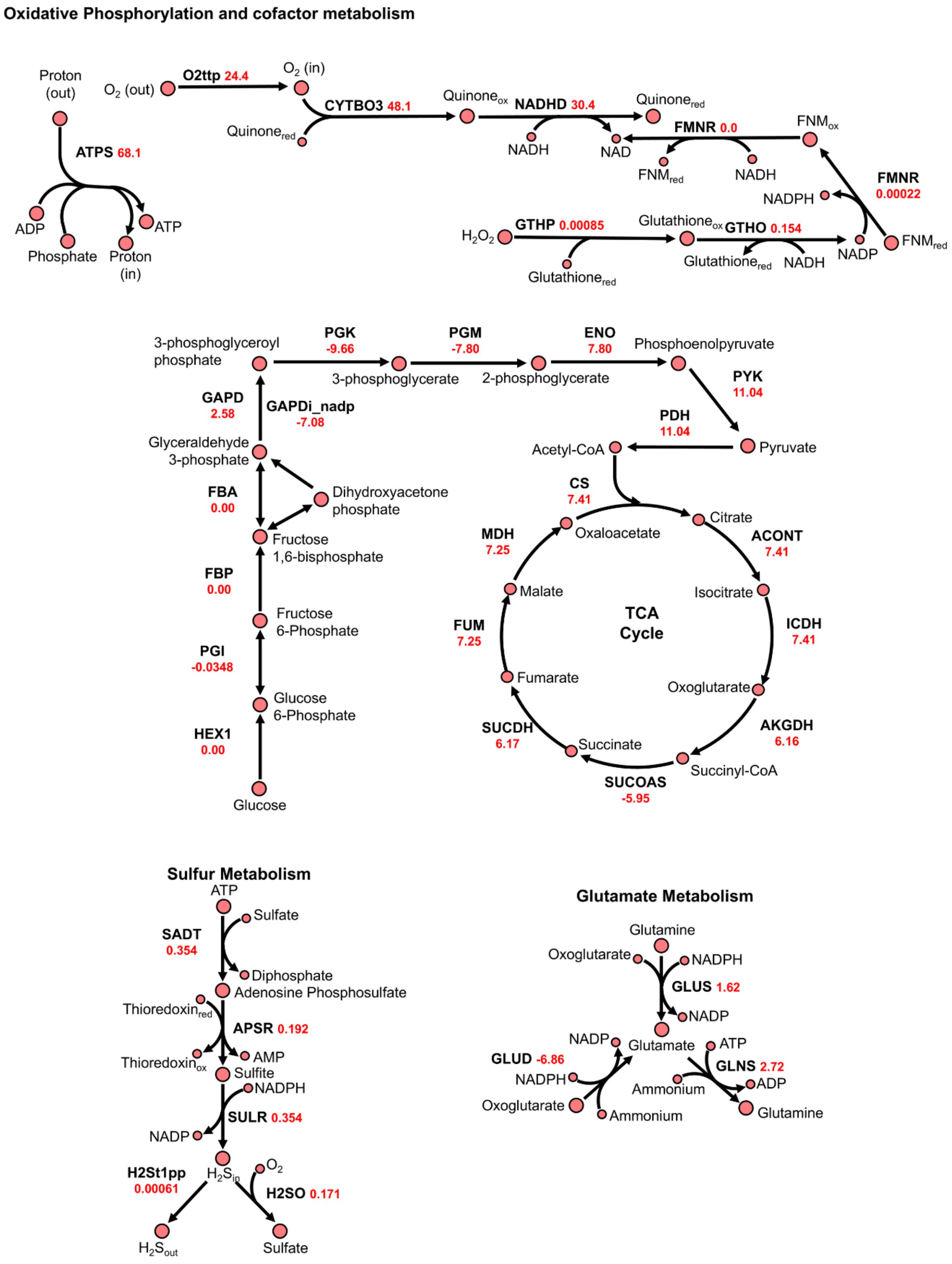


**Figure S6.** Pathway maps showing normalized fluxes for dextrose treatment.

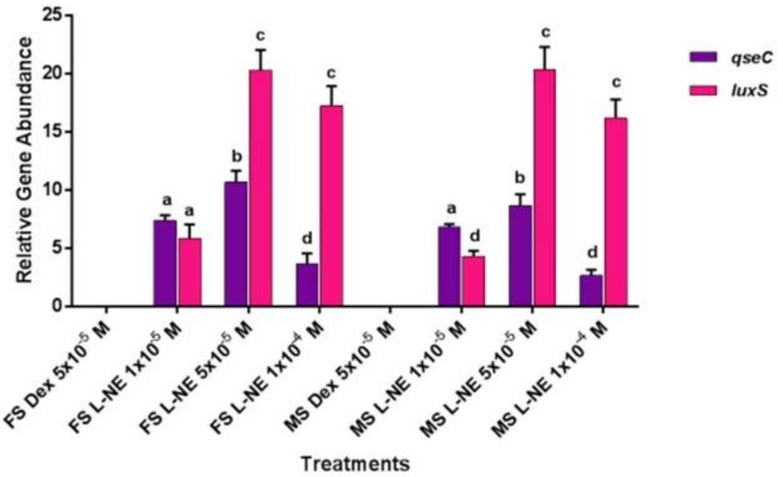


**Figure S7.** Relative Gene expression of *qseC* and *luxS* for both dextrose and L-NE at 24 hours. The initial inoculum of mixed bacteria for both media at time zero was 100 CFU/ml. Each point and bar represent the mean±SD of quadruplet (n=4) measurements. ($p<0.05$) denotes a significant difference within the dextrose (control) and L-NE treatment and between different time intervals.

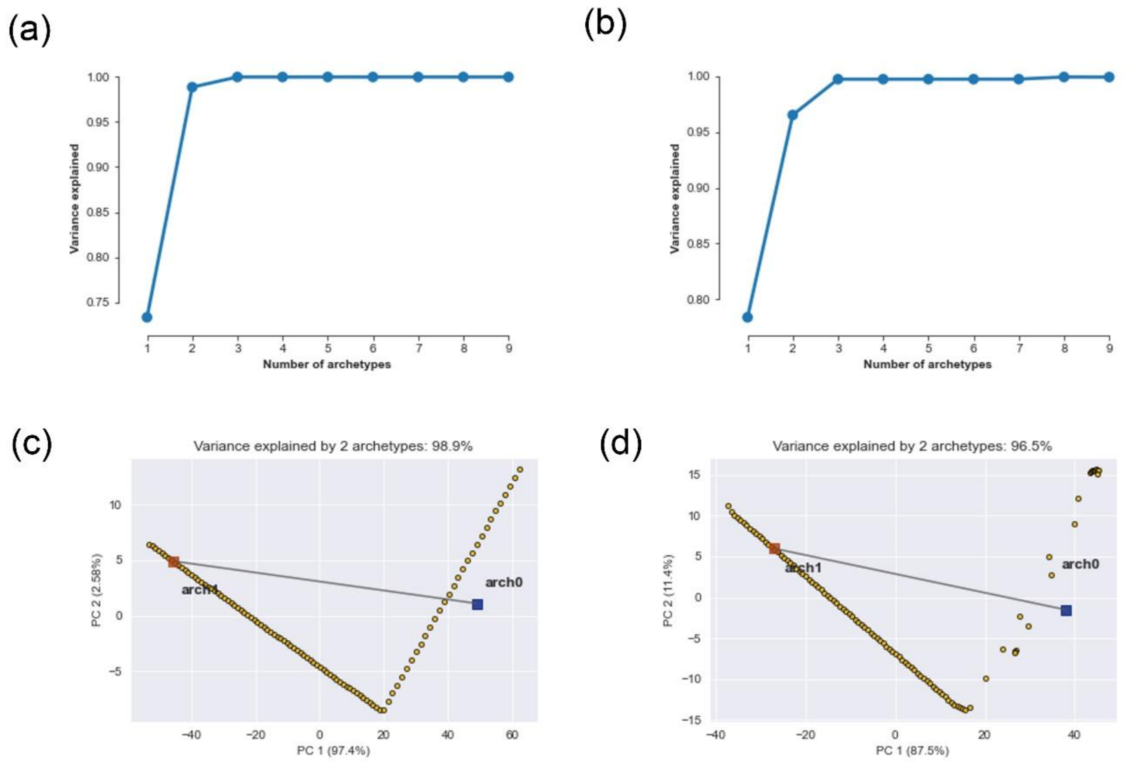


**Figure S8.** Two archetypes were computed for (a, c) dextrose and (b, d) L-NE conditions, respectively. From these, second archetype was chosen for further analyses because they corresponded to the set that reflected least deviation from the experimental data.

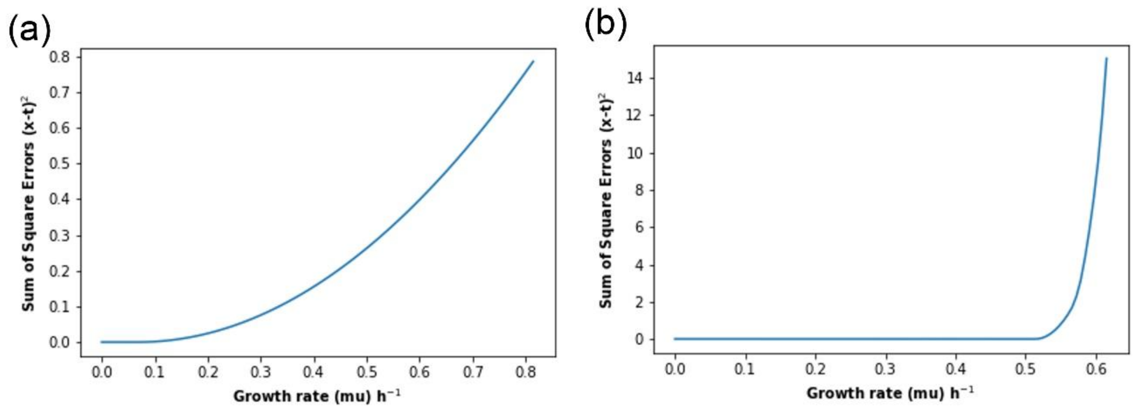


**Figure S9.** Using proteome expression data and model simulations, the fluxes that correspond to the metabolic state that represents the least deviation from the experimental data were computed. As can be seen in the figure, for (a) dextrose, the growth rate of <0.2 or lower reflects the lowest sum of squared errors, whereas for (b) L-NE, a growth rate of 0.5 or lower can be chosen.
